# Hierarchical cell-type identifier accurately distinguishes immune-cell subtypes enabling precise profiling of tissue microenvironment with single-cell RNA-sequencing

**DOI:** 10.1101/2022.07.27.501701

**Authors:** Joongho Lee, Minsoo Kim, Keunsoo Kang, Chul-Su Yang, Seokhyun Yoon

## Abstract

Single-cell RNA-seq enabled in-depth study on tissue micro-environment and immune-profiling, where a crucial step is to annotate cell identity. Immune cells play key roles in many diseases while their activities are hard to track due to diverse and highly variable nature. Existing cell-type identifiers had limited performance for this purpose. We present HiCAT, a hierarchical, marker-based cell-type identifier utilizing gene set analysis for statistical scoring for given markers. It features successive identification of major-type, minor-type and subsets utilizing subset markers structured in a three-level taxonomy tree. Comparison with manual annotation and pairwise match test showed HiCAT outperforms others in major- and minor-type identification. For subsets, we qualitatively evaluated marker expression profile demonstrating that HiCAT provide most clear immune cell landscape. HiCAT was also used for immune cell profiling in ulcerative colitis and discovered distinct features of the disease in macrophage and T cell subsets that could not be identified previously.

## Introduction

Since the first introduction of scRNA-seq technologies(1) and the subsequent elaboration(2,3), it has become one of the standardized methods for many biological researches. Unlike bulk RNA-seq, scRNA-seq enabled microscopic studies at single-cell resolution on intra- and inter-cellular activities in heterogenous tissue. Studies on tumor microenvironment and immune-profiling of many disease(4-7) are being actively conducted by many groups. An essential step to scRNA-seq analysis is cell-type identification(8,9) since a tissue is heterogeneous complex of cells with diverse phenotypic features(10) interacting with each other. Specifically, identification of immune cells and inferring their activation state are important issue in those studies. Due to its significance, there were vast amount of research for cell-type identification, either marker- or reference-based approaches. The former utilizes existing cell markers for given GEP. Garnett(11), SCINA(12), scSorter(13), scType(14), scCatch(15), DigitalCellSorter(16,17), CellAsign(18) and MarkerCount(19) are of this class, for which there are several databases usable for cell-type markers, such as Panglao DB(20) and CellMarker DB(21). While, the latter utilizes prior annotation to obtain cell-type profiles or markers to be used for identification, SingleR(22), scPred(23), scmap(24), CaSTLe(25), CHETAH(26), scHPL(27), clusstifyr(28), MARS (29), scPretain(30), Superscan, scLearn(31), scCapsNet(32), ACTINN(33), SciBet(34), scID(35), SingleCellNet(36), scVI(37), scMatch(38), scAnnotatR(39),. Although manual annotation using Seurat(40), SCRAN(41), SC3(42) or SCANPY(43) are also usable, it is time-consuming and not scalable for the explosive growth of scRNA-seq data generation. Among these cell-type identifiers, the majority was the reference-based(9). In comparisons of the two approaches(19) with the same datasets, the reference-based approaches were slightly better than the marker-based one. It can be due to the lack of good markers uniformly applicable to different tissues and/or its usage was not suitable. Compared to the marker-based approach, the reference-based identifiers search for data/tissue-specific markers from the prior annotation possibly leading to better results, even though one need to prepare the reference first.

Despite these tendencies, we focus on the marker-based approach since (1) it is easy to use requiring only markers. It can be a better option if we have good markers and utilize them appropriately. Then, how one can improve the marker-based approach? Our answer is to use the markers in a structured manner. There were studies on transcriptome-based cell-type taxonomy for scRNA-seq experiment(44,45). Based on transcriptomic profile, one can define different levels of cell-type taxonomies, for example, major-type, minor-type, and subset, for each of which one can devise identifiers separately and organize them in a hierarchy. In fact, distinguishing T cells and myeloid cells is not the same as classification of myeloid cell minor-types (e.g., distinguishing DC, monocyte, and macrophage). Classification of subsets, such as Tregs, Th1, 2, 9, 17, and 22, might be even different from minor-type classification since they typically represent different polarization states of the same minor type. Common markers in many subsets make subset identification harder, and one must not rely only on a few markers to annotate subset. For example, *SELL*(*CD62L*) is a common marker of monocyte, naïve CD4 T and Tregs. For such subsets, a better decision can be made if their minor-type is known *a priori*. As a matter of fact, the reference-based approach can hardly be used for identification of immune cell subsets simply because no references are available with ‘standardized’ subset nomenclature. There exist some datasets providing with subset annotations for specific subsets with well-recognized markers, such as Tregs or Th17. However, they are mostly non-standardized marked using only a few markers.

With these observations, we present HiCAT, a hierarchical, marker-based cell-type annotation tool, featuring improved performance for subset identification, particularly of immune cells. The method utilizes subset markers structured in a taxonomy tree. Although it roughly corresponds to the lineage tree, we rather define them by their transcriptomic profiles for classification purposes. There are previous works utilizing hierarchy in reference-based approaches(26,27), while we applied to marker-based method in a more structured way utilizing the taxonomy tree.

## Materials and Methods

### Datasets

For performance evaluation, we collected eight scRNA-seq datasets, including PBMC 68K (GSE93421) (46), CBMC 8K (GSE100866) (47), Melanoma 5K (GSE72056) (48), Lung 57K (ERP114453) (49), Lung 114K (SRP218543) (50), BALF 43K (GSE158055) (7), Colon 365K (SCP259) (5), and BRCA 100K (GSE176078) (6), where the first three were used many times in previous works for cell-type identification(19,26). Table 1 summarizes a brief specification of the datasets. Melanoma 5K and BRCA 100K contain tumor cell population, with which one can evaluate rejection function for unknown population. Most of all, the eight datasets contain lots of immune cells, including T cells, B cells and myeloid cells, which are our main focuses in the evaluation of the proposed method and other state of the art tools. Supplemental Figure 1 and 2 show UMAP plots for BRCA 100K and Colon 365K, and comparisons of the manual and HiCAT annotation as examples. For all the datasets, manual annotations are available for specific cell-type so that one can easily reannotate their minor and major-types for the purpose of performance evaluation in major- and minor-type level. Supplemental Figure 3 shows examples of reannotation of major- and minor-types given specific manual annotation.

**Table 1.** A summary of datasets used for the analysis and performance comparisons.

| Dataset | Accession | Protocol | Num. of genes | Num. of cells total | Num. of Tumor cells | Num of major /minor-types |
| --- | --- | --- | --- | --- | --- | --- |
| Colon 365K(5) | SCP259 | 10X genomics | 18,151 | 365,492 | None | 8/14 |
| BRCA 100K(6) | GSE176078 | 10X genomics | 29,733 | 100,064 | 24,489 | 8/17 |
| PBMC 68K(46) | GSE93421 | 10X genomics | 32,643 | 68,579 | None | 4/7 |
| CBMC 8K(47) | GSE100866 | Drop-seq | 20,501 | 8617 | None | 8/12 |
| Lung 57K(49) | ERP114453 | 10X v2 | 25,204 | 57,202 | None | 8/15 |
| Lung 114K(50) | SRP218543 | 10X v2 | 33,694 | 114,396 | None | 9/14 |
| BALF 43K*(7) | GSE158055 | 10X genomics | 27,943 | 42,723 | None | 6/11 |
| Melanoma 5K(48) | GSE72056 | Smart-seq2 | 23,684 | 4,513 | 1,251 | 6/8 |

\* Selected only bronchoalveolar lavage fluid (BALF) samples

### HiCAT processing

The brief procedure and the taxonomy tree used are depicted in Figure 1a and 1b, where the annotation is performed successively for major-type, minor-type and subsets along the tree. Although marker lists are defined only for subsets, those for minor-or major-types is obtained from subset markers by taking union for the subsets belonging to a specific minor-or major-type. The detailed procedure of HiCAT was depicted in Figure 1d, which was built using six elementary building blocks shown in Figure 1c. The building blocks includes (A) PCA and clustering, (B) Marker Counting and GSA scoring, (C) Unknown cluster detection, (D) GMM based correction, (E) Rejecting unclear cells, and (F) *k*NN based correction. Using these building blocks, HiCAT successively determines major-type, minor-type and subset, utilizing the subset markers structured in three-level taxonomy tree. The rationale is that one can possibly make better decision on subset identity knowing their broader type, especially when there exist common markers in several minor-types or subsets from different major-types.

**Figure 1.**
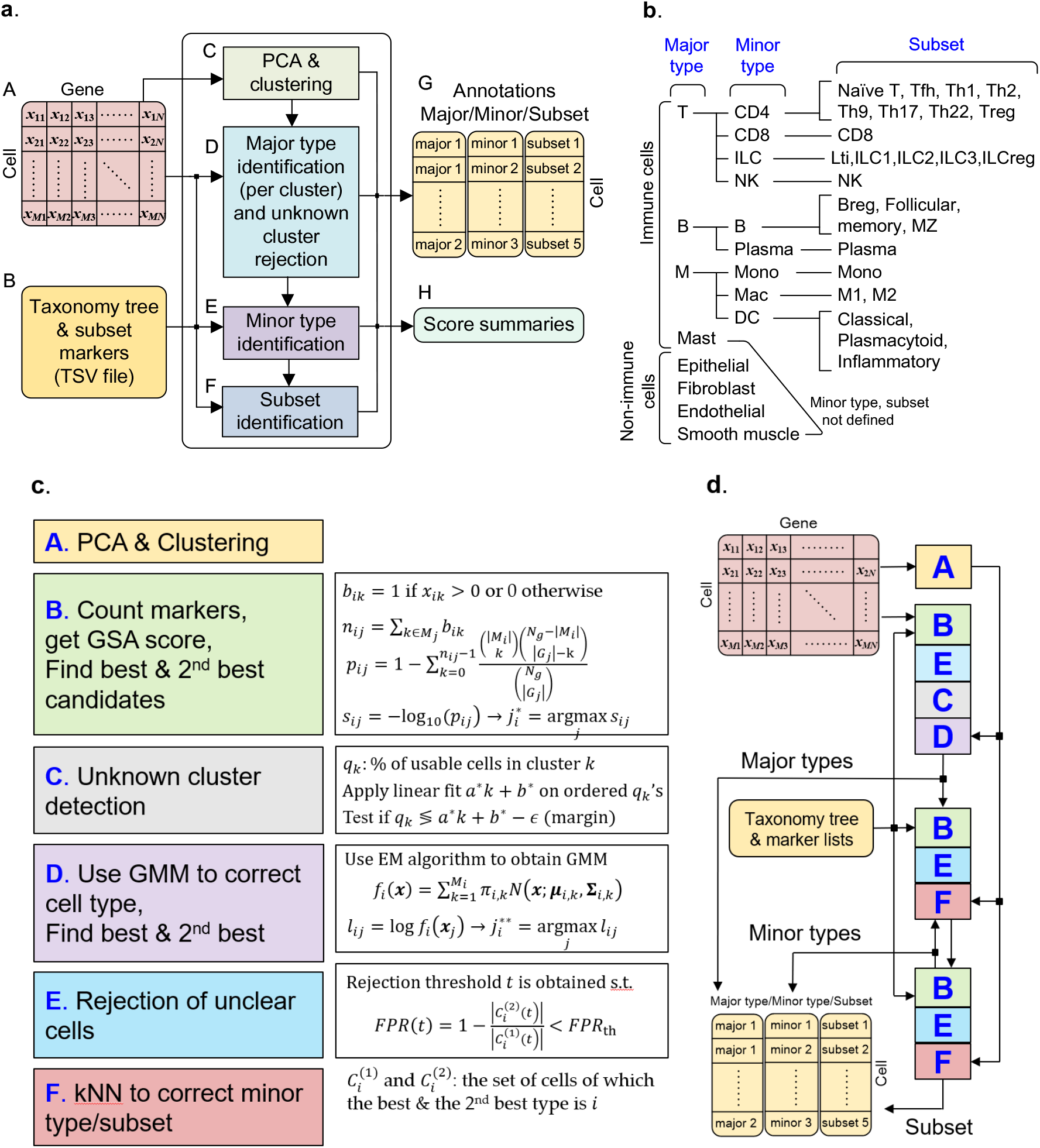
A brief HiCAT processing flow (a), taxonomy tree used for hierarchical identification (b), building blocks (c), and the detailed processing flow. In (a), the proposed identifier takes two inputs; a cell-by-gene matrix, A, and the marker DB, B. The cell-by-gene matrix is first PCA transformed and cells are clustered in dimension-reduced space, C. In major-type identification step, D, unclear clusters are rejected by using the best and 2^nd^ best major-type scores. Minor-type, E, and subset identification, F, follow to finally outputs 3-level annotation results, G, and score summaries, H. Note that markers are specified only for subsets. For major, the union of makers of its subsets was used, whereas, for minor-type, we used maximum of subset GSA scores within a minor-type. In the taxonomy tree (b), as our focus is on immune cell subsets, other non-immune cells were not further divided into minor-types or subsets. T: T cells, B: B cells, M: Myeloid cells, MZ: marginal zone B cell, Treg: regulatory T cells, Breg: regulatory B cells. Using the elementary blocks in (c), the HiCAT provide different processing for major-type, minor type and subset identification according to their respective transcriptomic features.

#### Three-level taxonomy tree

The taxonomy tree was shown in Figure 1b. Although the three levels of cell-type taxonomies roughly correspond to the lineage tree, we rather define them by their transcriptomic profiles for classification and clustering purposes. First, major-types are defined as cell populations that are clearly separable from others so that they are easily identifiable in a low-dimensional landscape, e.g., in tSNE or UMAP plots, while minor-types are a subpopulation of a major-type and not clearly separable even though they can be well localized within a major-type cluster. In our taxonomy, CD4, CD8 T cells, ILC and NK cells were set as minor-types of T cells. Subsets are a subpopulation of a minor-type and typically represents specific activation/polarization states of a minor population, e.g., M1 or M2 polarization of macrophage. They are not clearly separable from other subsets and also may not be well localized such that blind clustering is not applicable to catch a specific subset. As mentioned, these three-level taxonomies are defined only for the classification purposes and the development of hierarchical identifier. We implemented hierarchical identification based on the taxonomy tree in Figure 1b, where, since our primary interest was on immune cells, other non-immune cells were not divided into minor-types or subsets, and this can be considered as that there is only one minor-type or one subset.

#### Structured marker DB

To implement hierarchical identification, we need list of markers for each subset in the taxonomy tree, which are fed to the identifier as a tap-separated values file having 5 columns, including major-type, minor-type, subset, marker type and comma-separated list of markers (gene symbols). The first tap in Supplemental File 2 is the one we used in this work. The taxonomy tree can be easily recovered from the tuples of major-type, minor-type and subset. The marker type is one of positive (pos), negative (neg) or secretory (sec), which were adapted from the type of markers in R&D systems (https://www.rndsystems.com/resources/cell-markers) where we obtained the subset markers. Positive markers are either cell surface markers or intracellular markers that are typically expressed in a subset, regardless of its expression level. Negative markers are those not expressed. The negative markers are considered as a hit when evaluating GSA score if the cell does not express the negative markers. Although it provides useful information in minor-type or subset identification within a major-type (e.g., CD4+ and CD8+ T cell identification within a T cell cluster), it may not work in major-type identification if other major-types expresses those negative markers defined only for a few specific minor-type or subset. The last one is the secretory factors, of which the coded proteins are secreted by some subsets presenting their respective activation state. Although they are also an important information when identifying subsets, they are highly variable according to the biological state of the tissue they reside. Therefore, we used only positive markers in this work even though the usage of the 3 marker types can be user-specified. Using the subset marker defined in the marker DB, the markers of major-types can be defined as the union of markers for the subsets belonging to that major-type.

#### Input cell-by-gene matrix

Another input to the HiCAT is the cell-by-gene matrix, of which each row is a cell, and each column is a gene. The matrix should be formatted as a data frame, for which the row names are cell barcodes and column names are gene symbols as typical in a scRNA-seq data. The column names are used to identify the expression state of a specific marker.

#### Preprocessing

In the preprocessing step, we perform count normalization, log transformation and principal component analysis (PCA) by choosing highly variable genes, similar to those with Seurat(40) or SCANPY(43). Clustering is then performed to be used for the cluster-basis major-type identification and unknown cluster detection. HiCAT support three clustering algorithms, including *k*-means clustering, Louvain’s algorithm or GMM-based clustering. User can select either one even though we used Louvain’s algorithm in this work.

### Major-type identification and unknown cluster detection

This process is performed in 4 steps; (1) computing GSA scores, (2) rejection of unclear cells, (2) detection of “unknown cell-type” clusters, and (4) Gaussian mixture model (GMM) based correction.

#### Marker count and gene set analysis (GSA) scoring

For the first step in the cell-type identification, we need cell-type markers, where, for the major-types, they are given by the union of subset markers as mentioned before. Given the set of major-type markers, *M*_*i*_, of size |*M*_*i*_| for the *i*th cell-type and the gene expression matrix of size *N*_*c*_ × *N*_*g*_, with *N*_*c*_ being the number of cells and *N*_*g*_ the number of genes, the marker counts is given by *n*_*ij*_ = |*M*_*i*_ ∩ *G*_*j*_ | for the *i*th cell-type of the *j* th cell, where *G*_*j*_ is the set of genes expressed in the *j* th cell, regardless of their expression level. The GSA score is defined by

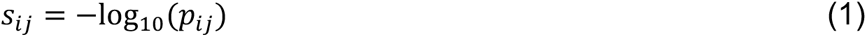

with the p-value, *p*_*ij*_, given by

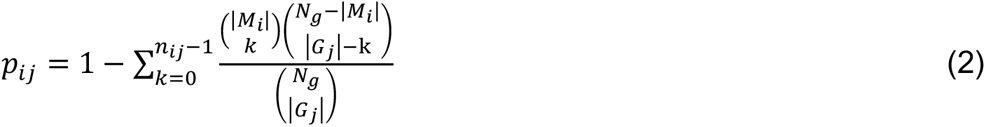

which is the probability of the marker counts *n*_*ij*_ is obtained by chance(51). Once the scores are obtained, the initial major-type of the *j*th cell is given by

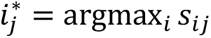

and its score is set to max_*i*_ *S*_*ij*_. But this process is performed in conjunction with the rejection of unclear cells in the next.

#### Rejection of unclear cells

Next step is to set threshold for rejection. Rejecting unclear cell is performed per-cell basis. Taking the batch effect into account, we need to use cell-type specific threshold that can be obtained from the dataset itself. To this end, we first determine the best and the 2^nd^ best type for each cell given *S*_*ij*_ for *i* ∈ *T, j* ∈ *C*, where *T* is the set of major-types to identify and *C* is the set of cells. Let 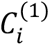 and 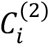 be the set of cells of which their best and the 2^nd^ best type is *i*. Apparently, they are disjoint to each other. As cells in 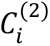 might be some other types than *i* with high probability, their score *S*_*ij*_ for *j* ∈ 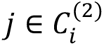 may give us a clue for the threshold of the *i*th cell-type.To be specific, define two set of cells for given rejection threshold 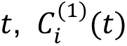 and 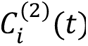 They are the set of cells from 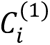 and 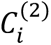, respectively, of which the GSA score is greater than the threshold *t*. We obtained the threshold *t*_*i*_ for the *i*th cell-type as the minimum of *t* satisfying the false positive rate defined below is less than a certain threshold, *FPR*_threshold_, i.e.,

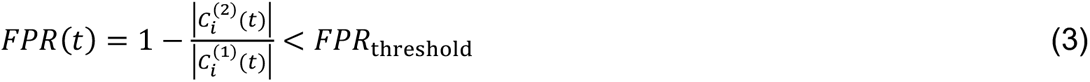

With the threshold *t*_*i*_ obtained, the cells with its score, max_*i*_ *S*_*ij*_ being less than *t*_*i*_ are replaced with ‘unassigned’.

#### Detecting unknown cell-type clusters

It is an important feature of cell-type identifiers to detect unclear cell-type cluster that can be possibly unidentified-so-far or tumor cells. The average score within a cluster can be used as an indicator of unknown cell-type. If there exist some unknown (major) type of cells that are not contained in the marker DB entries, the cell cluster might have much less scores than others. The problem is that it is hard to find a universal threshold that applies to every dataset as they might be obtained by different platforms utilizing different reference genomes and, sometimes, they were preprocessed in different ways. Moreover, different cell-types have different number of markers and different average scores. Therefore, it is not a good idea to use a universal threshold, but to use dataset specific threshold that can be obtained from the dataset itself. To identify those unknown clusters, we devised the following steps.

1. For each cluster *c*, the percentage, *q*_*c*_, of ‘usable’ cells with its score being greater than or equal to a certain threshold, *S*_DE_, is computed. Note that 0 < *q*_*c*_ ≤.
2. We sort *q*_*c*_‘s to obtain a sequence 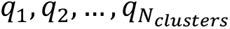, to which a linear fit is performed for high ranked *q*_*c*_‘s, where the fitting cost *J* is defined as the sum of squared error weighted by *q*_*k*_, i.e., 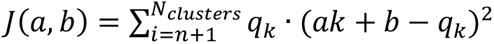, where *n* is given by a specific proportion of *N*_*clusters*_, e.g., round(*aN*_*clusters*_) with *α* =0.3.

(c) Once the fitted line *a*^∗^*k* + *b*^∗^ is obtained, the decision whether cluster *k* is kept or discarded is made by a test if *q*_*k*_ < *a*^∗^*k* + *b*^∗^ − *t*, where *t* = (1 − *α*) · max_*j*_ | *a*^∗^*j* + *b*^∗^ − *q*_*j*_ |. Supplemental Figure 4 shows example statistics for unknown cluster detection.

#### Gaussian mixture model (GMM) based correction

This is to reassign major-type to a cluster unintentionally excluded and or erroneously identified. Suppose that a cluster is marked ‘unknown’ even if it is not “clearly separable” from some nearby clusters with proper cell-type assignment. If it is not clearly separable from them, it might not be unknown, especially their major-type. To identify those clusters and reassign a major-type if applicable, we proceed as follows.

a. For each major-type, say the *i*th major-type identified in the previous steps, we model their distribution as a mixture of multivariate normal on the dimension reduced space, i.e.,

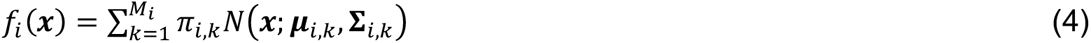

where ***π***_*i,k*_, ***μ***_*i,k*_ and **Σ**_*i,k*_ are the size, mean and covariance matrix, respectively, of the *k*th component of the *i*th major-type, *M*_*i*_ is the number of mixture components and ***x*** is the (PCA transformed) gene expression vector. Given the major-type assignment performed using GSA score, the model parameters can be obtained by using the EM algorithm, for which various software tools are available.
b. (b) With the GMM parameters for all the major-type, we compute the log-likelihood, *l*_*ij*_ = log*f*_*i*_(*x*_*j*_), for each cell using its feature vector *x*_*j*_ and the major-type are revised by finding the maximum of *l*_*ij*_, i.e., the major-type of the *j*th cell is given by argmax_*i*_ *l*_*ij*_ and its score by max_*i*_ *l*_*ij*_.
c. (c) *Cluster-basis major-type assignment*: Once unclear clusters were excluded, major cell-types are assigned per-cluster basis majority voting, i.e., for each cluster, simply the majority is identified and all the cells in that cluster are assigned by the same major-type regardless of their respective percentage.
d. (d) *Correction using aggregated network topology*: When using graph-based clustering, one can identify aggregated network of clusters using the interconnection strength between clusters. Based on this interconnection network of clusters, one can further identify tightly connected group of clusters to correct major-type identified in step (c). For example, let’s say 4 nearby clusters were tightly interconnected, where 3 of them were T cells, while one was unassigned (or B cell with relatively low average score). Then, the unassigned cluster is highly likely to be T cell and we can correct it to the majority in the tightly connected group. ‘Unassigned’ cell cluster rejected previously can be reassigned through this procedure.

Note that the last two steps are from the assumption that major cell-types are well clustered and clearly separable from other major-types and, therefore, not applicable to minor-type identification.

### Minor-type identification

Once major-type is assigned to all the clusters, minor-types are assigned separately to each major-type utilizing the taxonomy hierarchy. The assignment procedure is as follows.

a. For each major-type, say the *m*th major-type, select the relevant subsets and their markers.
b. Count expressed markers and compute GSA score, *S*_*ij*_ for *j* ∈ *C*_M_ *i* ∈ *S*_M_, in the same way as for major-type assignment, where *C*_M_ is the set of cells assigned to the *m*th major-type and *S*_M_ is the set of subsets belonging to the *m*th major-type.
c. Minor type score is the maximum subset score among those of subsets belonging to that minor type, i.e., 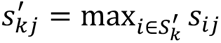 where 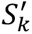 is the set of subsets belonging to the *k*th minor type.
d. Once the minor type scores are obtained, the initial minor-type of the *j* th cell is given by 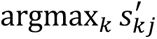 and its score is set to 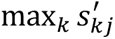.
e. *Application of k nearest neighbor (kNN) rule for minor-type correction*: Since minor-types are assumed to be roughly clustered, but not clearly separable to each other, we apply *k*NN rule to correct minor-type in PCA transformed space, i.e., for each cell in *C*_M_, select *k* nearest neighbors and take majority among *k* to correct its minor-type. By doing so, each minor-type can be roughly divided within the clusters of the corresponding major-type. For minor type correction, we set *k* to 31 in all experiments, even though it can be specified by the user.

### Subset identification and minor-type correction

Finally, subset assignment is performed similar to the minor-type assignment. Although *k*NN-rule based correction is supported for subsets as well, we set *k* = 1 for kNN rule (no correction) since there is no evidence reported so far if subsets are well localized within minor-type cluster.

### Running other marker-based methods

We compared HiCAT with 6 existing marker-based methods, Garnett(11), SCINA(12), scSorter(13), scType(14), scCatch(15), DigitalCellSorter(16,17). For CellAssign, we couldn’t install the package due to error. Although some of them provide their own list of markers, we used R&D systems markers (https://www.rndsystems.com/resources/cell-markers) in all methods for fair comparison and their default parameter settings. The usage of each method was reproduced from their Github page, except for Garnett and scSorter. We referred https://cole-trapnell-lab.github.io/garnett/ for Garnett and https://cran.r-project.org/web/packages/scSorter/vignettes/scSorter.html for scSorter. All the existing marker-based methods were run 3 times, each for major-type, minor-type and subset identification, respectively, while HiCAT was run once for each dataset since it provides the 3 levels of cell-types simultaneously. Since some of identifiers could not be run for big-sized datasets, such as BRCA 100K, Lung 114K, and Colon 365K, we ran all the identifiers separately for pair of samples.

### Performance criteria

#### Match test with manual annotation

To evaluate performance for major- and minor-type, we used five performance measures used in previous works(19,26), i.e., correct (C), error (E), erroneously assigned (EA), correctly unassigned (CUA), and erroneously unassigned (EUA). To define these criteria, we divide the cells based on their prediction results into ‘unassigned’ and others with valid predicted label (the cell-types existing in the marker DB). The unassigned is counted as CUA if its original label (the manual annotation) is not in the cell type list in the marker DB. Otherwise, they are counted as EUA. A cell with valid predicted label is counted as EA if its original label is not in the cell type list in the marker DB, or, counted as C if its original label exists in the cell type list and is equal to the predicted label. Otherwise, it is counted as E, i.e., if its original label exists in the cell type list but is not equal to the predicted label. The graphical explanation was depicted as reference in Figure 2 and 3.

**Figure 2.**
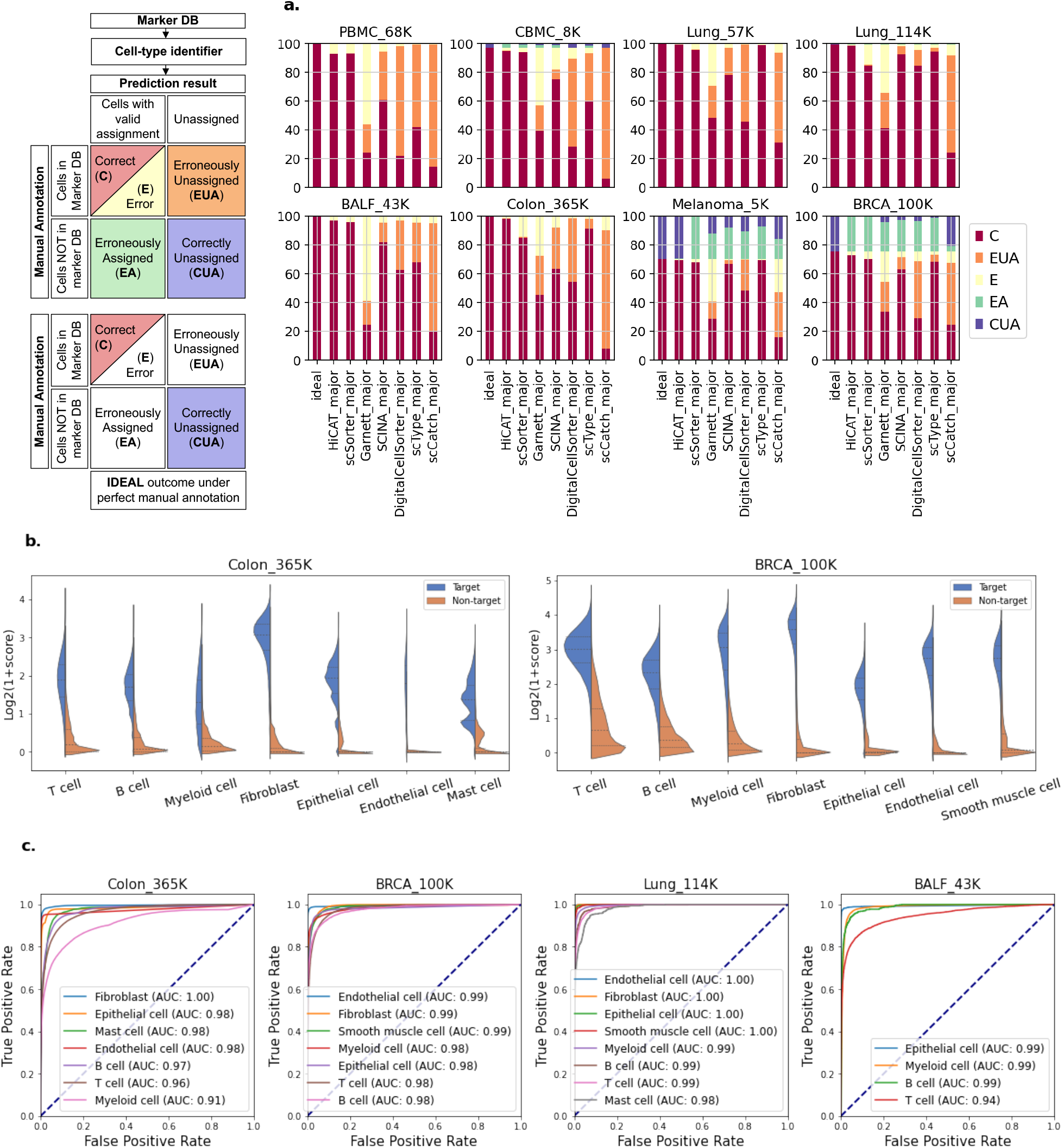
Comparisons of major-type identification performances (a), major-type GSA scores distribution of target cell-type versus non-target for Colon 365K and BRCA 100K (b), and ROC curves before and after GMM-based correction for the two datasets (c). In (a), the first bar marked as ‘ideal’ shows the ideal performance, where CUA account for tumor cells or those cell-types not in the marker DB entry, which should ideally be ‘unassigned’ as briefly represented on the right. Excluding CUA portion in the ideal performance, HiCAT showed 95% or higher accuracy in most datasets, except for PBMC 68K. In (b), GSA score distribution for target is the distribution of ‘target’ cell-type scores for the cells identified correctly as ‘target’, i.e., for those cells of which the cell-type with the highest score was the ‘target’, while that for non-target is the score distribution of the ‘target’ for the cells of which the 2^nd^ best cell type was the ‘target’, i.e., the best one was something else. The violin plots in (b) show that major-types are largely separable from others using only the GSA scores, even though there are small overlaps between the scores of targets and non-targets. Rejection thresholds for each major-type are set using the two distributions such that false positive rate is less than or equal to a pre-defined value, say 0.05. ROC curves in (c) confirm the separability of most major-types.

**Figure 3.**
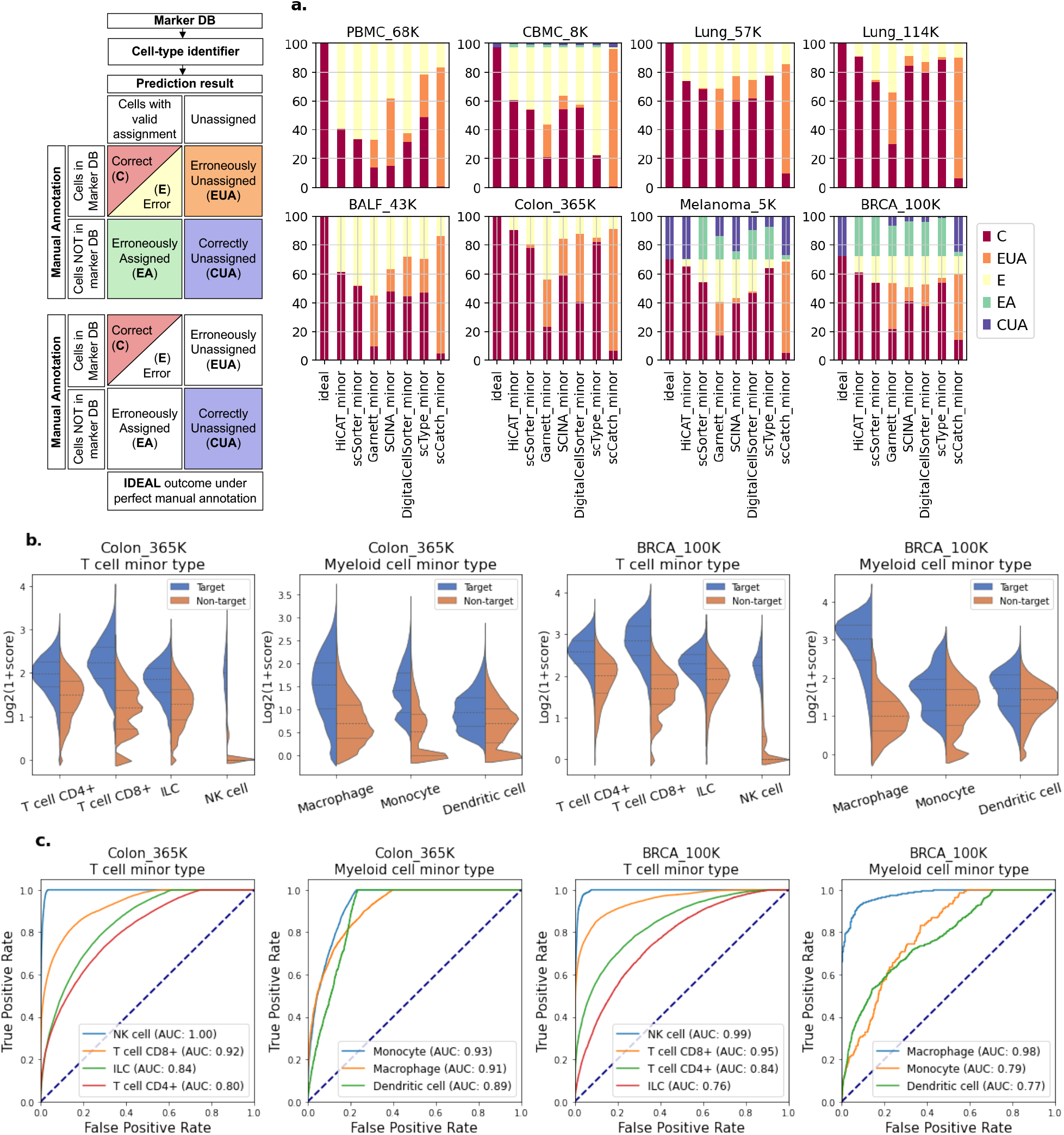
Comparisons of minor-type identification performances (a), T cell and myeloid cell minor-type GSA scores distribution for Colon 365K and BRCA 100K (b), and ROC curves for T cell and myeloid cell minor-type identification for the two datasets (c). In (a), the first bar marked as ‘ideal’ shows the ideal performance, where CUA account for tumor cells or those cell-types not in the marker DB entry, which should ideally be ‘unassigned’. Identification performances for minor-type are much less than those for major-types in all identifiers raising question whether manual annotations are reliable enough to be used as ground truth for performance evaluation. In (b), comparing with those for major-types, there exist considerable overlap between the two distributions for target and non-target, respectively, indicating that they may not be well separable using only the GSA scores. In minor-type identification, instead of cluster-basis reassignment, we used *k*-nearest neighbor (*k*NN) rules to correct initial decisions. The number *k* can be pre-specified by the user and we set it 31 in all experiments in this work. In (c), the AUCs are shown to be much smaller than those for major-types, reflecting minor-type identification is more complicated.

#### Pairwise match test

In pairwise match test, instead of using the 5 criteria below, we simply compared the two results from two different identifiers, where we counted the fraction of cells for which the two results are equal, regardless of whether they are ‘unassigned’ or valid cell-types, i.e., it is counted as a hit even if the two results are both ‘unassigned’.

### DEG and GSE analysis

Differentially expressed gene (DEG) and gene set enrichment (GSE) analysis were performed for immune cell profiling in ulcerative colitis. Seurat package was used for DEG and FGSEA package for GSE analysis. The analyses were performed separately for the two datasets, Colon 365K and Colon 32K. First, DEG analysis was performed in inflamed (test) versus healthy (control) condition setting, separately for all myeloid cell subsets and T cell subsets. Then, DEGs were obtained with adjusted p-value cutoff 0.05. GSE analysis was then performed using DEGs. Additionally, DEGs for M2a/M2b/M2c versus M1 was obtained to run GSE analysis to check possible M1-skewed polarization of the three M2 macrophage subtypes.

## Results

The identification performances were evaluated for major-type, minor-type and subsets, separately. For major- and minor-type identification, we used the ‘manual’ annotations provided by the data publishers as the ground truth, while, for subset identification, we used pairwise match test and qualitative comparison of marker expression profiles under annotations using various marker-based identification tools.

### Major-type identification performance

#### HiCAT shows the best performance in most datasets

Figure 2a shows comparisons of major-type identification performance with the existing marker-based identifiers for 8 datasets with manual annotation available, where 2 datasets (BRCA 100K and Melanoma 5K) contain tumor cells. In Figure 2a, we used five performance criteria used in the previous works(19,26), i.e., correct (C), error (E), correctly-unassigned (CUA), erroneously-assigned (EA), erroneously-unassigned (EUA). HiCAT was the best in terms of correct criterion, except for PBMC 68K, for which scSorter was slightly better than HiCAT. Considering only valid major-types, i.e., not including tumor cells, HiCAT showed mostly greater than 95% accuracy, except for PBMC 68K and BRCA 100K, where, in BRCA 100K dataset, the error was mostly for cancer epithelial cells that most of the identifier classified as (normal) epithelial cells. The major-types contained in the marker DB include T cell, B cell, myeloid cell, epithelial cell, fibroblast, endothelial cell, smooth muscle cell and mast cell. These cells are typically well clustered with sufficient separation from others. Even with these characteristics, some identifiers didn’t perform good enough, depending on datasets. For datasets with tumors, HiCAT almost perfectly identified tumor cells in Melanoma 5K by correctly unassigning any labels, while, in BRCA 100K, most of the identifiers, including HiCAT, annotated them as normal epithelial cell, except for scCatch, which, however, the performance was the worst.

#### Primary statistics and ROC/AUC show good separability of major-types

To give insight into how HiCAT works, we showed in Figure 2b and 2c the distribution of major-type GSA scores along with ROC and AUC. Figure 2b shows that major-types are largely identifiable using their scores while there are small overlaps between the two distributions of GSA scores for the designated cells and the non-designated, respectively. The overlap between the two distribution indicates the harness of the identification, reflecting possible identification errors when using only GSA scores. The errors can be corrected based on their similarity in GEP (proximity in feature space), for example, by using the GMM-based corrections, which will be discussed in the Online Method section. GMM identifies the region of population for each major-type and, by using the likelihood for given GEP, one can improve the performance as shown by the ROC/AUC before and after the GMM-based correction. ROC/AUC also indicate how well they are separable from others.

### Minor-type identification

#### Performance for minor-types were worse than those for major-types in all identifiers raising question whether manual annotation of minor-types is reliable enough

Compared to major-types, minor-types are not well clustered so that the performance is expected to be worse than those for major-types. Figure 3 shows similar plots to Figure 2, where, as expected, the overall performances are much lower than those for major-type. Comparing identifiers, HiCAT was the best in six datasets while scType was the best in two datasets, PBMC 68K and Lung 57K. Comparing datasets, PBMC 68K was the worst, for which all identifiers showed less than 50% of correct assignment. CBMC 8K also couldn’t be well predicted by most identifiers. This result raises a question whether manual annotation is reliable enough to be used as ground truth for performance evaluation, especially for minor-types and subsets. In Figure 2a and 3(a), the performance was evaluated assuming the manual annotations are perfect. It is plausible assumption for major-types since major-types are well clustered with enough separation between them so that one can easily determine major-types for each separable cluster by checking expression of a few well-known major-type markers, for example *CD3* for T cells, *CD79B* for B cells. Unfortunately, however, it is not the case for minor-types as well as subsets. In many cases, minor-types of a major-type do not clearly separable even though they may form a cluster localized within a larger cluster of a major-type (Supplemental Figure 1 and 2). Manual annotation is typically done in cluster-basis, i.e., all cells within a cluster (obtained blindly by a clustering algorithm) are assigned by the same cell-type so that the annotation precision is limited by clustering resolution. In major-type annotation, it isn’t a big problem since they are largely separable from others. While, in minor-type annotation, a cluster can be spanning over two or more minor-types, with which annotation error may occur.

#### Pairwise match test shows higher match rate than those with manual annotation and HiCAT showed the best match

Accounting possibly erroneous annotation in minor-types, it would be interesting to check how well prediction results from different identifiers match to each other. Surprisingly, some identifier pairs had better match than pairs with manual annotation. In Table 2(a) and (b), we summarized the pairwise prediction match for major- and minor-type identification where we showed identifier pair and their match rate for the best, 2^nd^ best and the best with manual annotation. First of all, the average of the best match rates with manual annotation (the last column) in Table 2(b) was 72.3% across all datasets, while the one for the best pairs was 85.6%. Of note, in most datasets, HiCAT was one of the best pairs, except for CBMC 8K, for which scSorter-DigitalCellSorter pair showed 85.4%, slightly higher than the second-best match rate 85.0% with HiCAT-DigitalCellSorter pair. The higher match rate can happen if specific processes and/or algorithms used in two identifiers in a pair are similar to each other so that they produce similar bias and outcomes, regardless of their actual precision. If it is the case, there must exist one majority of pairs in the best pair list. In Table 2(a) and (b), however, there exist different pairs, three HiCAT– scType, two HiCAT–scSorter, two HiCAT–Manual annotation and one scSorter–DigitalCellSorter pair. Although HiCAT was shown in the best pairs for seven datasets, it is paired with different identifiers and, therefore, the argument above cannot be justified. Even for the best match with manual annotation, HiCAT appeared 6 times among 8. The best match rate with manual annotation for major-type in Table 2(a) was higher than 94.1% while it was 72.3 for minor-type. This reflects not only the difficulty of identifying minor-types compared to major-type, but also that the manual annotation of minor-type may contain errors to some extent. Therefore, it is risky to use them as ground truth for performance evaluation and/or as references for reference-based identifiers. The difficulty of minor-type identification is also clear from their GSA score distributions. Figure 3b and 3c show the minor-type score distribution and their ROC and AUC for Colon 365K and BRCA 100K, respectively. Compared to those for major-type, two distributions for the designated (target) and the non-designated (non-target) considerably overlap and the AUCs are mostly smaller than those in Figure 2c.

**Table 2.** A summary of pairwise prediction match for (a) major-type and (b) minor-type identification, showing the best match, 2^nd^ best match and the best match with manual annotations, and (c) for subset identification, showing the best, 2^nd^ best and 3^rd^ best match. Tumor cells are considered to be ‘unknown’ and counted as match if they were ‘unassigned’ by two identifiers in a pair. In (b), the average best match rate (85.6%) is higher than the best match rate with manual annotation (72.3%), raising question whether manual annotation is reliable enough for performance evaluation against it. Of note, in both (b) and (c), HiCAT was one of the best match pair in most datasets, i.e., seven out of eight.

| **a.** | Dataset | Best match pair | Best match rate | 2nd Best match pair | 2nd Best match rate | Best tool with manual annot. | Best match rate with manual annot. |
| --- | --- | --- | --- | --- | --- | --- | --- |
| **Major-type** | PBMC_68K | HiCAT - scSorter | 95.2 | Manual - scSorter | 92.9 | scSorter | 92.9 |
|  | CBMC_8K | HiCAT - scSorter | 95.4 | HiCAT - Manual | 95.3 | HiCAT | 95.3 |
|  | Lung_57K | HiCAT - Manual | 99 | Manual - scType | 98.7 | HiCAT | 99 |
|  | Lung_114K | HiCAT - Manual | 98.6 | HiCAT - scType | 95 | HiCAT | 98.6 |
|  | BALF_43K | HiCAT - scSorter | 97.5 | HiCAT - Manual | 96.7 | HiCAT | 96.7 |
|  | Colon_365K | HiCAT - Manual | 98.5 | HiCAT - scType | 92.1 | HiCAT | 98.5 |
|  | Melanoma_5K | HiCAT - Manual | 98.9 | scSorter - SCINA | 82.6 | HiCAT | 98.9 |
|  | BRCA_100K | HiCAT - scType | 92 | HiCAT - scSorter | 91.5 | HiCAT | 73.1 |
|  | Average rate |  | 96.9 |  | 93.1 |  | 94.1 |
| **b.** | Dataset | Best match pair | Best match rate | 2nd Best match pair | 2nd Best match rate | Best tool with manual annot. | Best match rate with manual annot. |
| **Minor-type** | PBMC_68K | HiCAT - scSorter | 78.5 | scSorter - DCS | 57 | scType | 48.8 |
|  | CBMC_8K | scSorter - DCS | 85.4 | HiCAT - DCS | 85 | HiCAT | 61 |
|  | Lung_57K | HiCAT - scType | 85.8 | Manual - scType | 77.1 | scType | 77.1 |
|  | Lung_114K | HiCAT - scType | 88.2 | HiCAT - SCINA | 85.5 | HiCAT | 84.9 |
|  | BALF_43K | HiCAT - scSorter | 79.1 | HiCAT - SCINA | 69 | HiCAT | 61.4 |
|  | Colon_365K | HiCAT - Manual | 90.4 | HiCAT - scType | 84 | HiCAT | 90.4 |
|  | Melanoma_5K | HiCAT - Manual | 94.9 | Manual - scType | 71.3 | HiCAT | 94.9 |
|  | BRCA_100K | HiCAT - scType | 82.7 | HiCAT - scSorter | 76.6 | HiCAT | 60.1 |
|  | Average rate |  | 85.6 |  | 75.7 |  | 72.3 |
| **c.** | Dataset | Best match pair | Best match rate | 2nd Best match pair | 2nd Best match rate | 3rd Best match pair | 3rd Best match rate |
| **Subset** | PBMC_68K | HiCAT - scSorter | 60.3 | HiCAT - scType | 58.1 | scSorter - scType | 50 |
|  | CBMC_8K | HiCAT - scType | 67.8 | HiCAT - scSorter | 66.4 | scSorter - scType | 66.1 |
|  | Lung_57K | HiCAT - scType | 66.2 | scSorter - scType | 65 | HiCAT - scSorter | 61.7 |
|  | Lung_114K | HiCAT - scType | 63.9 | SCINA - scType | 56.1 | HiCAT - SCINA | 55.8 |
|  | BALF_43K | Garnett - scCatch | 52 | HiCAT - scSorter | 41.4 | HiCAT - scType | 41.3 |
|  | Colon_365K | HiCAT - scType | 77.7 | HiCAT - scSorter | 64.8 | scSorter - scType | 59 |
|  | Melanoma_5K | HiCAT - scType | 54.1 | scSorter - scType | 51.8 | Garnett - scCatch | 51.1 |
|  | BRCA_100K | HiCAT - scType | 74.2 | HiCAT - scSorter | 70.3 | scSorter - scType | 68.8 |
|  | Average rate |  | 64.5 |  | 59.2 |  | 56.7 |

\* DCS: DigitalCellSorter

### Subset identification

Identification of subsets, for example, M1 and M2a/b/c/d polarization of macrophages or Th1, Th2 and Th17 in CD4+ T cells, is even more challenging since they mostly represent activation states of the same minor-type and, sometimes, the states are reversible. Although some datasets provide subset annotation for immune cell subsets using well-recognized markers, they are mostly non-canonical so that they are not reusable for various downstream analysis other than those in the original publication and, most of all, it is impossible to evaluate performance for subset. Therefore, pairwise match rates were first evaluated as did for minor-type.

#### HiCAT-scType pair showed the best match in most datasets

Table 2(c) summarizes the best, 2^nd^- and 3^rd^-best match pair and their respective match rates. The average match rate over all datasets were 64.5, 59.2 and 56.7 for the best, 2^nd^- and 3^rd^-best pair, respectively, which are all even lower than those for minor-type identification, reflecting it is much more difficult to precisely identify cell-type subsets. Regardless of this tendency, HiCAT-scType pair showed the best match rate in 6 datasets among 8. As mentioned before, it may be because similar processing and algorithms were used in the two identifiers. However, from the results for minor-type identification, it seems not the case. Without the ground truth available, we cannot say which one is better than the other, even though HiCAT was better in minor-type identification.

#### Qualitative evaluation of marker expression profile

Although quantitative evaluation is impossible, we can give insight into HiCAT performance by showing marker expression profile for each identified subset. Figure 4 shows a landscape of marker expression profile under the annotations with different identifiers, including HiCAT, scSorter, SCINA and DigitalCellSorter, for Colon 365K. The figure shows at a glance how well the annotated cell type(subsets) represent the expressed markers in their designated subset population. This type of analysis is used in manual annotation to determine cell-type for each cluster. Here, we used it to check how well the identifier determined cell-type subsets. Ideally, the marker genes, except for common markers, must be expressed only in their designated subsets but not in the non-designated subsets, even though cell type subsets cannot be clearly defined. However, since they represent specific activation states of the same cells, we need to consider intermediate states between them. In the figure, we need to check two things, the percentage of cells in a subset that are expressing the designated marker genes (dot size) and their mean expression levels (color). For better visualization, we ordered the marker genes according to the dot size for each subset.

**Figure 4.**
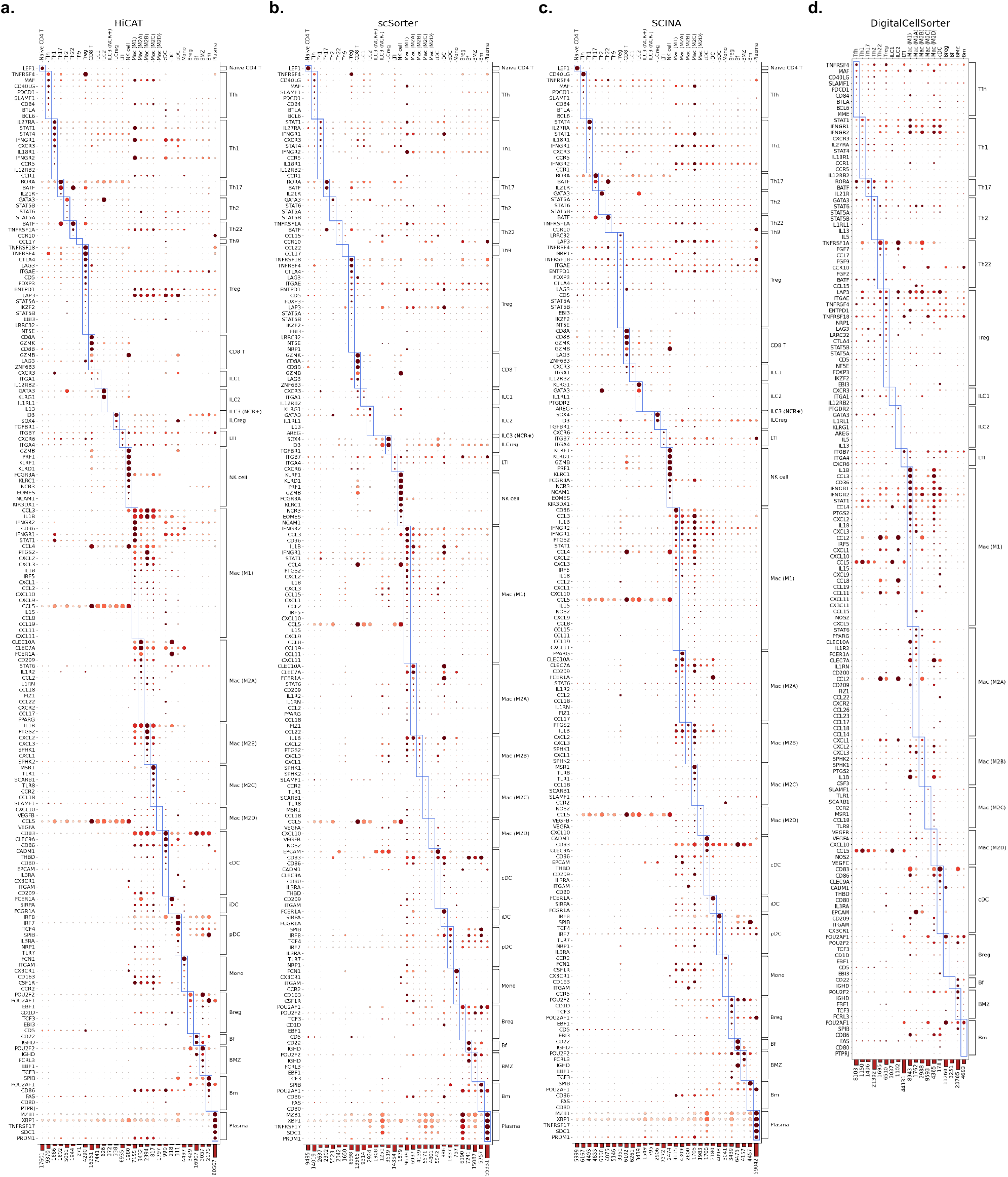
Dot plots showing marker expression profiles of immune cell subsets in Colon 365K under annotations from different tools, (a) HiCAT, (b) scSorter, (c) SCINA, and (d) DigitalCellSorter. The dot size is the fraction of cells, in an identified subset, that expresses the marker, regardless of their expression level. The color corresponds to the expression level scaled between 0 and 1, where 0 and 1 represent respectively the minimum and the maximum expression level of that marker among all subsets. The fraction was clipped at 0.6, i.e., fractions greater than 0.6 were set to 0.6. For better visualization, we grouped the marker genes for each subset and ordered according to the fraction of marker expressing cells in the target subsets. Each column is a subset identified by each method. HiCAT, scSorter, and SCINA identified 29/30 immune cell subsets, while DigitalCellSorter identified only 19 subsets, respectively. We showed up to 180 markers with highest percentage of marker expressing cells.

#### HiCAT annotation shows the best marker expression profile in immune cell subsets

Figure 5a shows the percentage of marker expressing cells (MEC) of four datasets for up to 200 markers represented as a curve, where the markers were ordered according to their percentages. Comparing the identifiers that we examined, HiCAT showed higher percentage of marker expressing cells for most of the markers. In Figure 5b, we chose only T cell subsets in Colon 365K and showed their marker expression profiles, up to 80 high percentage markers, for the four identifiers that showed highest percentage curves. Rows of a dot plot are the subsets identified by the corresponding tool so that, if there are subsets not showing in the rows, it means the identifier decided those subsets not existing in the datasets. HiCAT, scSorter, and SCINA identified all 9 T cell subsets, while scType identified only 4 (other identifiers found 6 or less even though we did not show them here). Although one can argue that there indeed exist only 4 T cell subsets, it is not likely because other 3 tools identified all 9 T cell subsets. Note that one must not assess the quality of annotation relying only on a few markers or separately for each subset. Many markers must be considered for the qualitative evaluation. Overall, the results in Figure 5 implicitly indicate better identification of immune cell subsets by using HiCAT.

**Figure 5.**
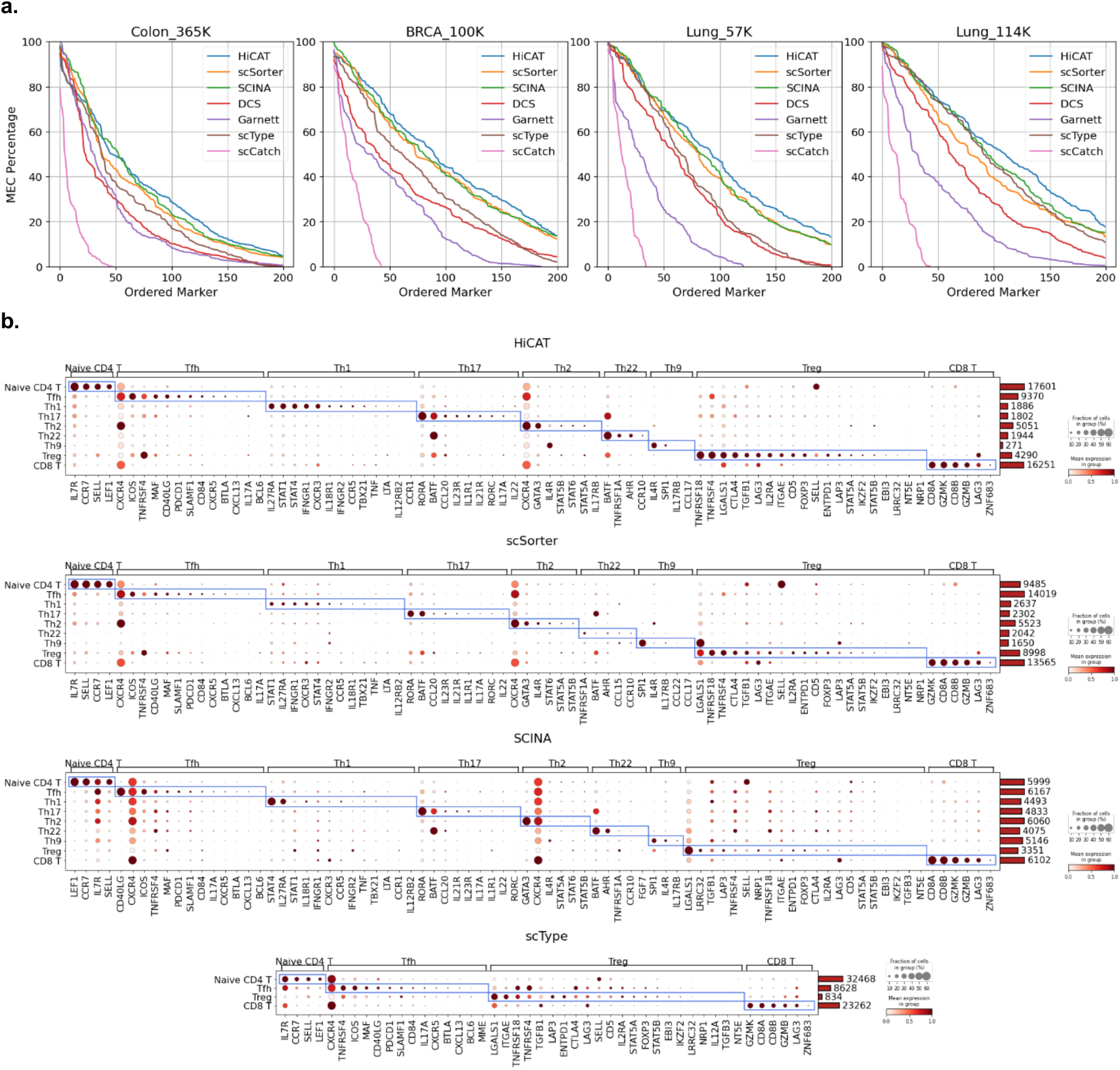
A comparison of the percentage of marker expressing cells (MEC) for four datasets (a) and dot plots showing marker expression profile of T cell subsets in Colon 365K under annotations from different tools (b). In (a), we showed the percentage of MEC for up to 200 marker genes, reversely ordered according to its MEC percentage. HiCAT showed higher MEC percentage for most of the markers. Although scSorter and SCINA showed higher percentage than HiCAT for some markers, they are decreasing faster than HiCAT. In (b), we showed the marker expression profiles of T cell subsets under annotations from the four identifiers with higher percentage curves of MECs in (a). Each row is T cell subset identified by each method. HiCAT, SCINA and scSorter identified all 9 T cell subsets, while scType identified only 4 subsets, respectively. We showed up to 80 markers with highest percentage of marker expressing cells.

The fractions of marker expressing cells in its designated subset were mostly much larger than those in non-designated subset, even though some of the markers appeared not or rarely expressed. Some markers have non-negligible fraction in non-designated subsets, e.g., *CXCR4* for Tfh and Th2. Their mean expression levels, however, are less than those in the designated subset. *BATF* is common markers of Th17 and Th22 so that they were shown to be comparable in all their designated subsets.

### Immune cell profiling in ulcerative colitis using HiCAT

Macrophages are phagocytes in tissues and play a crucial role in intestinal mucosal homeostasis, regulation of inflammatory response, and protection against pathogens(52,53). Macrophages generally consist of M1 and M2 macrophages, where the M2 macrophages are further classified into M2a, M2b, M2c, and M2d subtypes. M1 macrophages produce pro-inflammatory cytokines such as IL-6, IL-1β, and TNF-α, whereas the M2a, M2b, and M2c subtypes are associated with the responses to anti-inflammatory activity(54,55). However, an aberrant polarization of macrophages such as abundant M1 and depleted M2 could be observed in the progression of UC(56) and M1-skewed transcriptional program in macrophages were known to ignite chronic activation of NLRP3 inflammasome causing inflammatory bowel disease (IBD), such as ulcerative colitis(UC)(57-59). Although M2 macrophages are associated with anti-inflammatory activities in general, M2b is known to secret pro-inflammatory cytokines as well, including IL-6, IL-1β, and TNF-α, acting both pro- and anti-inflammatory reaction, and play an important role in the IBD(60).

In this subsection, HiCAT is used for high resolution immune profiling in ulcerative colitis, which may provide us a deeper insight into the complicated disease. A previous study profiled more than 365,000 single-cell transcriptomes (Colon 365K) and provided a comprehensive overview of epithelial, stromal, and immune cell subsets associated with UC patients at the single-cell level(5). However, immune cell subsets were not thoroughly identified in the study, possibly due to the lack of accurate cell type identification tools for minor cell types. In this dataset, samples were collected from two groups, heathy(*n*=12) and colitis patients(*n*=36). The samples from the latter were further divided into ‘inflamed’ (*n*=18) and ‘non-inflamed’ (*n*=18). Here, we use the dataset to provide in-depth profiling of immune cells, focusing on macrophage and T cell subsets, in terms of canonical nomenclature. To confirm the results from Colon 365K, we additionally used another dataset, GSE182270 (Colon 32K), where two groups of samples are provided (4 healthy and 5 inflamed). Since the number of samples in Colon 32K is only 9, we used it only for supplementary to the result from Colon 365K.

### M2b macrophages are overpopulated in inflamed condition, while M2a is underpopulated

We have identified three myeloid lineages, dendritic cells, monocytes and macrophages, and also their subsets (Figure 6a). Since macrophages are involved in the pathology of ulcerative colitis(52), we compared the composition of five macrophage subsets (M1, M2a, M2b, M2c, and M2d) in each sample (Figure 6b). Interestingly, M2b macrophages in inflamed condition were significantly higher in percent than those in healthy while M2a were reversed (Figure 6c). This finding was also supported by the other dataset, Colon 32K, showing the same tendency for M2a and M2b, even though M1 and M2c were shown to be significantly different as well. Since the number of samples is limited in Colon 32K, we accept the results from Colon 365K. The marker expression profiles of macrophage subsets in UC and healthy condition (Figure 6d) shows that common markers of M1 and M2b, including *IL1B, CCL3, PTGS2, CXCL2*, and *CXCL3*, are stronger in M2b than in M1 under the inflamed condition, possibly reflecting M2b is more active and has stronger proinflammatory signals.

**Figure 6.**
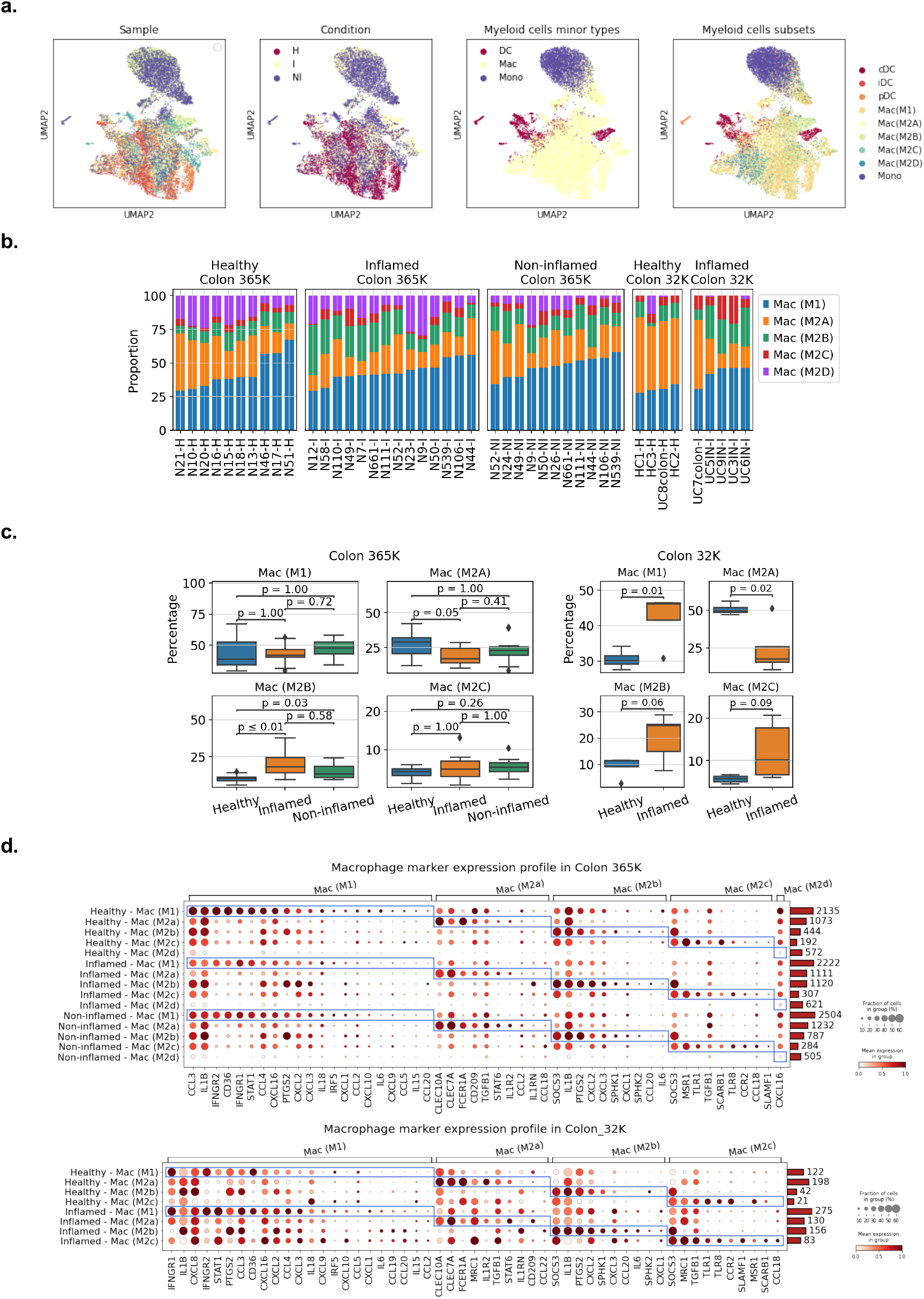
Single-cell transcriptomic landscape of macrophage subsets in Colon 365K and Colon 32K. (a) UMAP plots for Colon 365K, (b) relative population of macrophage subsets in different conditions in Colon 365K and Colon 32K dataset, (c) Box plots for the percentage of macrophage subsets in different conditions and the statistical significance of population differences between two conditions, and (d) the marker expression profiles of macrophage subsets in different inflammatory conditions. In (b) and (c), we see that, in both datasets, M2b was shown to be overpopulated in inflamed condition, comparing with healthy condition, whereas M2a was underpopulated in inflamed. Although M1 macrophage was shown to be overpopulated in inflamed condition in Colon 32K dataset, it couldn’t be identified in Colon 365K.

### M1-skewed M2b macrophage matters in ulcerative colitis

To get more insight into their modulated polarization, DEG and GSE analysis were performed using Seurat and FGSEA(61), respectively. The results were attached in Supplementary File 2. We found from the DEG and GSE analysis for inflamed versus healthy that M2b in inflamed condition shows higher enrichment scores in four pathways, including TNF□ signaling via NF□B, signaling by interleukins, inflammatory response, and cytokine signaling in immune system, with adjusted p-value less than 0.05. Comparing M2b and M1 in inflamed condition, these pathways were also identified and had also higher enrichment scores with the same p-value threshold. Since these pathways were not identified in other test-control pairs, the result confirms that M2b plays critical role in the chronic inflammation of ulcerative colitis. This result could be observed in Colon 32K as well. Overall, the reanalysis of two independent scRNA-seq datasets of ulcerative colitis using HiCAT clearly showed that inflamed sites in UC patients had an imbalance of macrophage subsets (high M2b and low M2a) compared to healthy control samples, suggesting that M2b among M2 macrophages might be responsible for the inflammatory responses observed in UC patients.

### Overpopulation of Treg and underpopulation of Th1 and Th2 in inflamed condition

Composition of T cell subsets were also significantly different in inflamed condition from those in healthy (Figure 7). In Colon 365K dataset, Treg, Th17 and Th22 were shown to have significantly higher proportion in inflamed condition than in healthy, while Th1 and Th2 were reversed. In Colon 32K, Treg, Th1 and Th2 showed similar tendency, even though they were not significant. Nevertheless, overpopulation of Treg and underpopulation of Th1 and Th2 seem to be general tendencies in inflamed condition. The results for DEG and GSE analysis on each CD4+ T cell subsets showed that inflammation related pathways in Tfh and Th2 were deactivated (Supplemental File 2).

**Figure 7.**
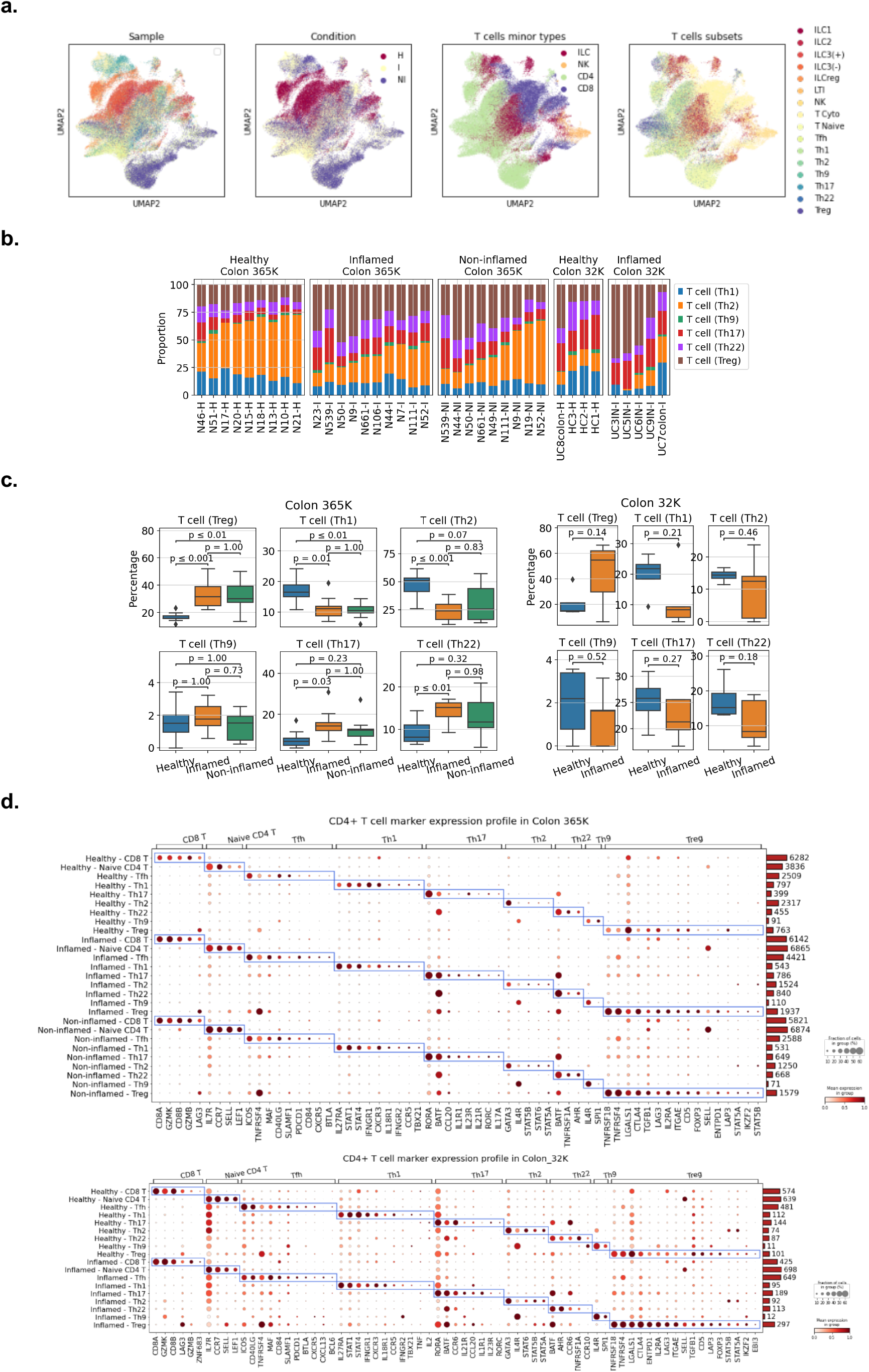
Single-cell transcriptomic landscape of T cell subsets in Colon 365K and Colon 32K. (a) UMAP plots for Colon 365K, (b) relative population of T cell subsets in different conditions in Colon 365K and Colon 32K dataset, (c) Box plots for the percentage of T cell subsets in different conditions and the statistical significance of population differences between two conditions, and (d) the marker expression profiles of T cell subsets in different inflammatory conditions. In (b) and (c), comparing the inflamed to the healthy condition, Treg was overpopulated in the inflamed condition, whereas Th1 and Th2 were reversed. These tendencies were shown to be statistically significant in both datasets. Th17 and Th22 were shown to be significantly overpopulated in the inflamed condition in Colon 365K, while they were not confirmed in the other dataset.

## Discussion

There are many huddles in cell-type annotation in single-cell RNA-seq data, especially for annotation of minor types or subsets. Here, we presented a hierarchical cell-type identifier, HiCAT, implemented in 3-steps using cell-type taxonomy tree and subset marker database. Performances were evaluated quantitatively for major and minor types and qualitatively for subset. For major type identification, HiCAT showed the best match rate against manual annotation with mostly higher than 95% except for PBMC 68K. For minor type identification, it showed the best match with manual annotation in 6 datasets among 8. Considering possible annotation error in minor type manual annotation, we also evaluated pairwise match among all identifiers including manual annotation. In this experiment, HiCAT was appeared to be one of the best match pair for most datasets, except for CBMC 8K. This result indicates that HiCAT is more reliable than others including manual annotation. For subset identification, no dataset provides subset annotations in canonical nomenclature and, hence, we could only evaluate the pairwise match, where HiCAT-scType pair was the best in 6 datasets. The comparison of marker expression profiles for immune cell subsets showed HiCAT was the best in the clarity of subset marker expression pattern. Finally, the HiCAT annotation for Colon 365K revealed distinct composition of macrophage and CD4+ T cell subsets. Providing clearer transcriptomic landscape of the disease, we could find that M1-skewed M2b macrophage is responsible for the chronic inflammation in ulcerative colitis. Certainly, these findings could not be obtained without precise annotation of immune cell subsets using HiCAT.

The successive identification possibly gives us a better result in minor-type and subset identification. Specifically, immune cell subsets typically do not have clear boundaries between them. Rather, they form “a continuum beginning with hard and ending with soft distinctions among cell-types with an ambiguous transition between these extremes” (45). In this case, simultaneous identification of all minor-types or a mix of minor-types and subsets will not give us a good result. HiCAT seems to work well by successively determining cell’s identity given their broader class as a condition for statistical scoring of lower classes.

Another aspect of HiCAT is that cluster-basis annotation is used only for major-type identification while *k*-nearest neighbor (*k*NN) rule is used for minor-types and subsets. In many identifier, cell-types are assigned in cluster-basis, i.e., the same cell-type is assigned to all cells in a cluster. In this case, annotation accuracy is limited by clustering resolution and it is desired to perform clustering with high resolution setting. However, if it is too high, the statistics on marker expression become inaccurate as the number of cells in a cluster gets smaller. On the other hand, if the resolution is too small, two or more subsets may be grouped into one cluster leading to incorrect assignment. Therefore, cluster-basis assignment is not a good option for subset annotation, while it is fine for major or possibly some minor-types. HiCAT applies cluster-basis assignment only for major-type. While, in minor-type and subset, the annotation is initially made based only on the GSA score. Cell-type correction is then applied using *k*NN, with different number *k* for minor-type and subset identification. Effectively, the *k*-nearest-cells form a micro cluster, in which the center cell’s identity is determined (corrected) by majority voting

Another thing to account when using markers for cell-type annotation is the fact that transcript expression does not match to the protein expression. CD4 and CD8 surface markers in T cells are always present on their surface so that one can make sure they are CD4+ or CD8+ using antibodies for physical cell sorting. However, this does not hold when counting RNAs since they are not necessarily present in their transcriptome at the time of measurement. Therefore, annotation using only a few markers defined in protein level is risky to use confidently. If a minor-type is well clustered, one can identify them cluster-by-cluster basis since some portion of the cells in the cluster is expressing the markers in transcriptome level. However, this may not be the case for subsets, and it is desirable to use as many markers as possible simultaneously to make better decisions using their statistics. From these unique features, HiCAT could reliably be used for identification of immune cell subsets. Although we cannot say the subset annotation of HiCAT is completely reliable, one can select confident cells using GSA scores provided in the HiCAT output.

## Abbreviations

AUC: Area under ROC curve
DC: dendritic cell
DB: database
DEG: Differentially expressed gene
EM: Expectation maximization
GEP: gene expression profile
GMM: Gaussian mixture model
GSA: Gene set analysis
GSE: Gene set enrichment
IBD: inflammatory bowel disease
*k*NN: *k*-Nearest Neighbor
ROC: Receiver operating characteristic
scRNA-seq: single-cell RNA-seq
tSNE: t-distributed stochastic neighbor embedding
UMAP: Uniform Manifold Approximation and Projection
Th: T helper cell
Treg: regulatory T cell
UC: ulcerative colitis.

## Data and Software Availability

The python code and the example in Jupyter notebook were deposited to PyPI and Github, respectively. Installation instruction, usage and example codes can be found at https://github.com/combio-dku/HiCAT (Project name: HiCAT, license: GPL 3.0, Operating system(s): Platform independent, Programming language: python 3, other requirement: None). The datasets used in this work can be freely downloaded from the internet: two lung datasets from the human cell atlas, https://www.humancellatlas.org/, and others from the gene expression omnibus (GEO) at https://www.ncbi.nlm.nih.gov/geo/ using their accession number.

## Authors’ contribution

SY, KK and CY devised the concept and key idea. SY developed the code of HiCAT. JL and MK performed experiments. SY and KK guided experiments and data analysis. KK gave interpretation and feedback on the performance evaluation. CY curated immune profiling of ulcerative colitis. All authors wrote, read, and approved the final manuscript.

