## Supplemental Fig for "Hierarchical cell-type identifier accurately distinguishes immune-cell subtypes enabling precise profiling of tissue microenvironment with single-cell RNA-sequencing"

### ABSTRACT

Single-cell RNA-seq enabled in-depth study on tissue micro-environment and immune-profiling, where a crucial step is to annotate cell identity. Immune cells play key roles in many diseases while their activities are hard to track due to diverse and highly variable nature. Existing cell-type identifiers had limited performance for this purpose. We present HiCAT, a hierarchical, marker-based cell-type identifier utilizing gene set analysis for statistical scoring for given markers. It features successive identification of major-type, minor-type and subsets utilizing subset markers structured in a three-level taxonomy tree. Comparison with manual annotation and pairwise match test showed HiCAT outperforms others in major- and minor-type identification. For subsets, we qualitatively evaluated marker expression profile demonstrating that HiCAT provide most clear immune cell landscape. HiCAT was also used for immune cell profiling in ulcerative colitis and discovered distinct features of the disease in macrophage and T cell subsets that could not be identified previously.

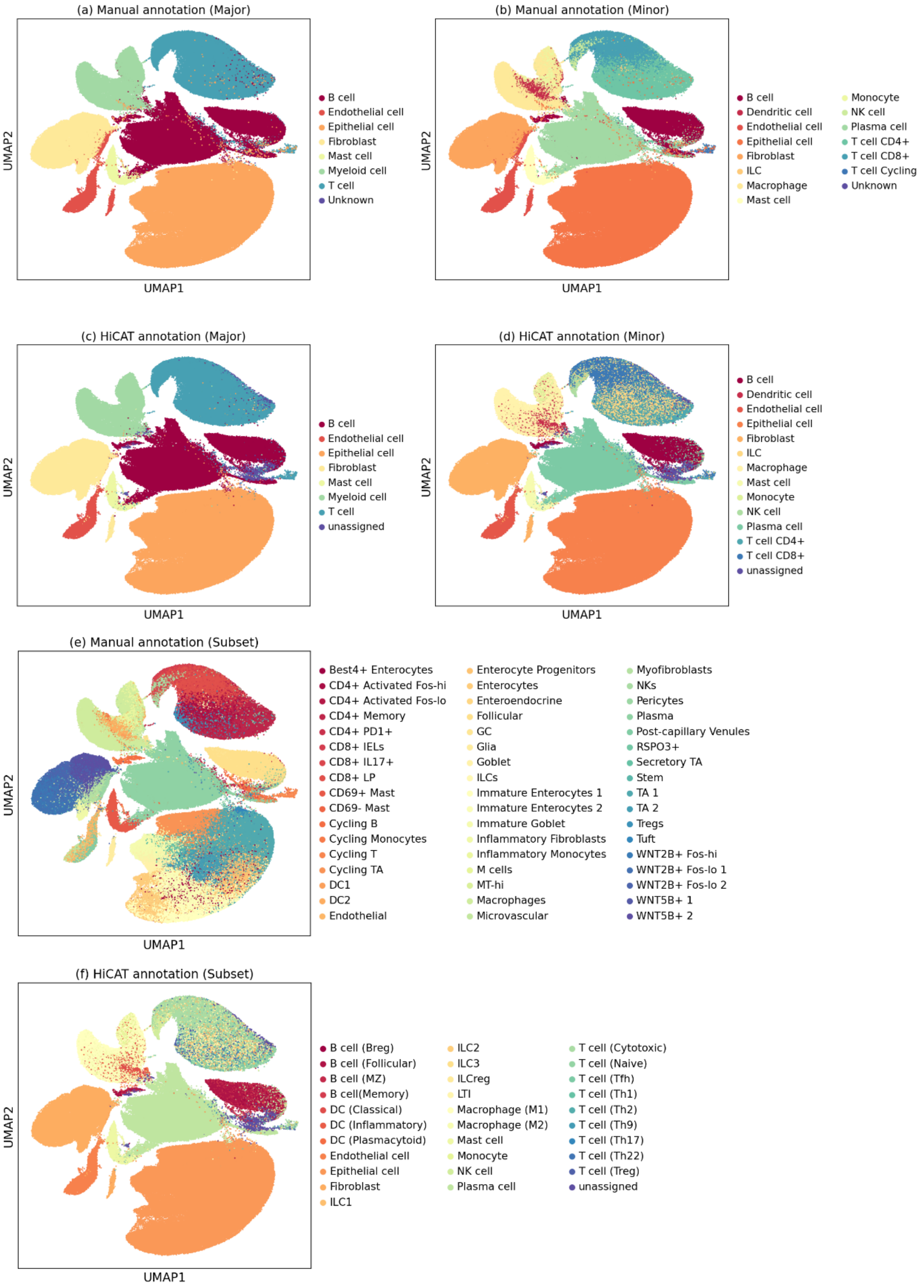


**Supplemental Figure 1.** UMAP plot and comparison of manual annotation and HiCAT results for Colon 365K datasets. (a), (b), and (e) are manual annotation of major-type, minor-type and subset (non-canonical), respectively. (c), (d), and (f) are HiCAT annotation of major-type, minor-type and subsets (canonical), respectively. Comparing (a) and (b), major-types are clearly separable from others with sufficient separation between them, while minor-types, for example T cell and myeloid cell minor types, are not clearly separable even though they are mostly well localized within a major-type cluster. Nevertheless, it is questionable if manual annotation of minor types is reliable enough to evaluate identifier performance against it.

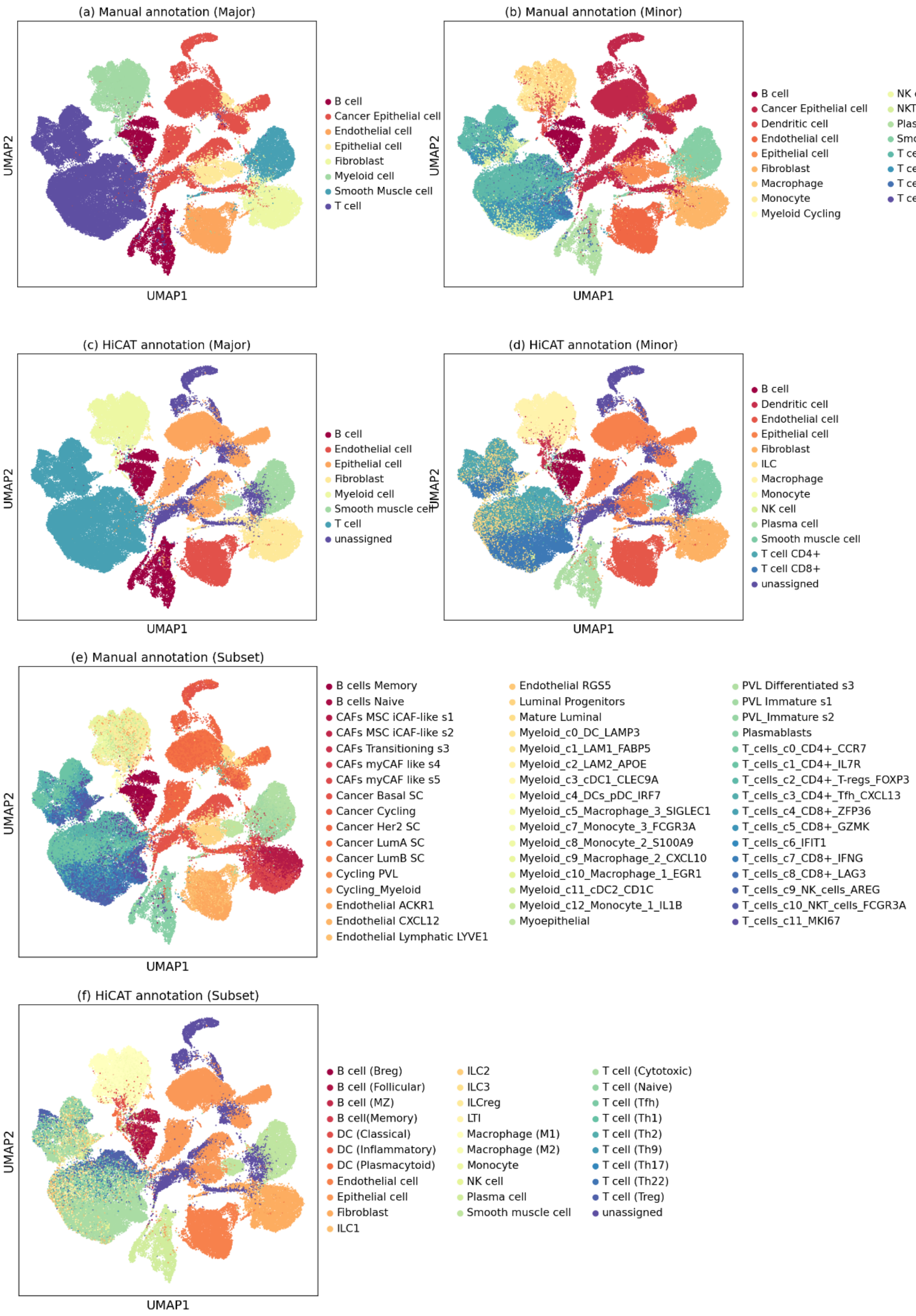


**Supplemental Figure 2.** UMAP plot and comparison of manual annotation and HiCAT results for BRCA 100K datasets. (a), (b), and (e) are manual annotation of major-type, minor-type and subset (non-canonical), respectively. (c), (d), and (f) are HiCAT annotation of major-type, minor-type and subsets (canonical), respectively. Like Supplemental Figure 10, major-types in (a) are clearly separable from others, while minor-types in (b), for example T cell and myeloid cell minor types, are not clearly separable even though they are mostly well localized within a major-type cluster.

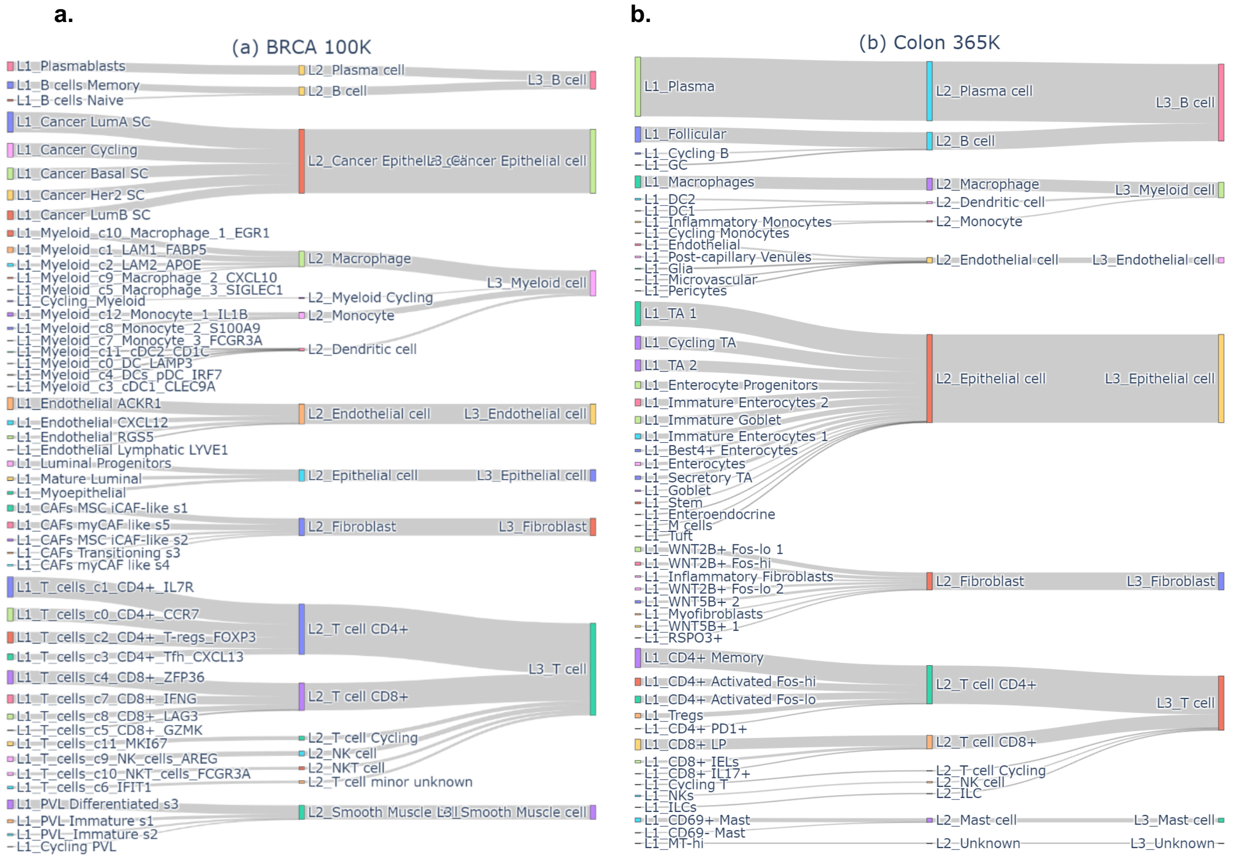


**Supplemental Figure 3.** Examples of setting minor-type and major-type given the specific type annotation contained in the datasets for BRCA 100K (a) and Colon 365K (b). Left: manual annotations provided along with the dataset, Center: reannotated minor-types, Right: reannotated major-types. We used these minor-types and major-types to compared with the decisions of HiCAT and other existing identifiers, excluding 'Unknown's and cycling cells.

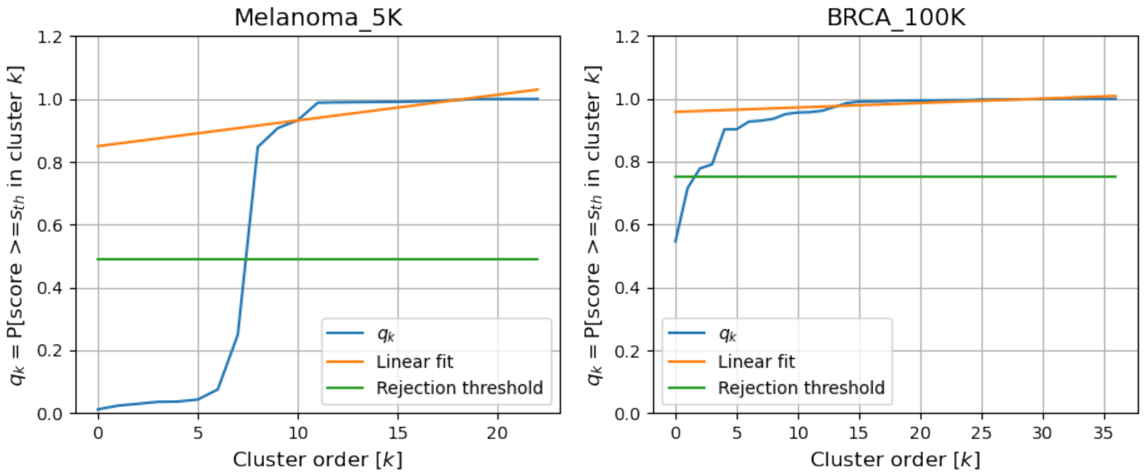


**Supplemental Figure 4.** Statistics for unknown cluster rejection for (a) Melanoma 5K and (b) BRCA 100K. The cluster index is ordered according to the percentage of 'usable' cells, $q_k$, defined by $Pr\{\text{GSA score} \geq \text{threshold}\}$ for cluster $k$. Linear fitting was applied with those $q_k$'s for upper 60% (reconfigurable) and the rejection threshold is obtained from the maximum difference between the fitted line and those $q_k$'s for lower 40% of clusters.

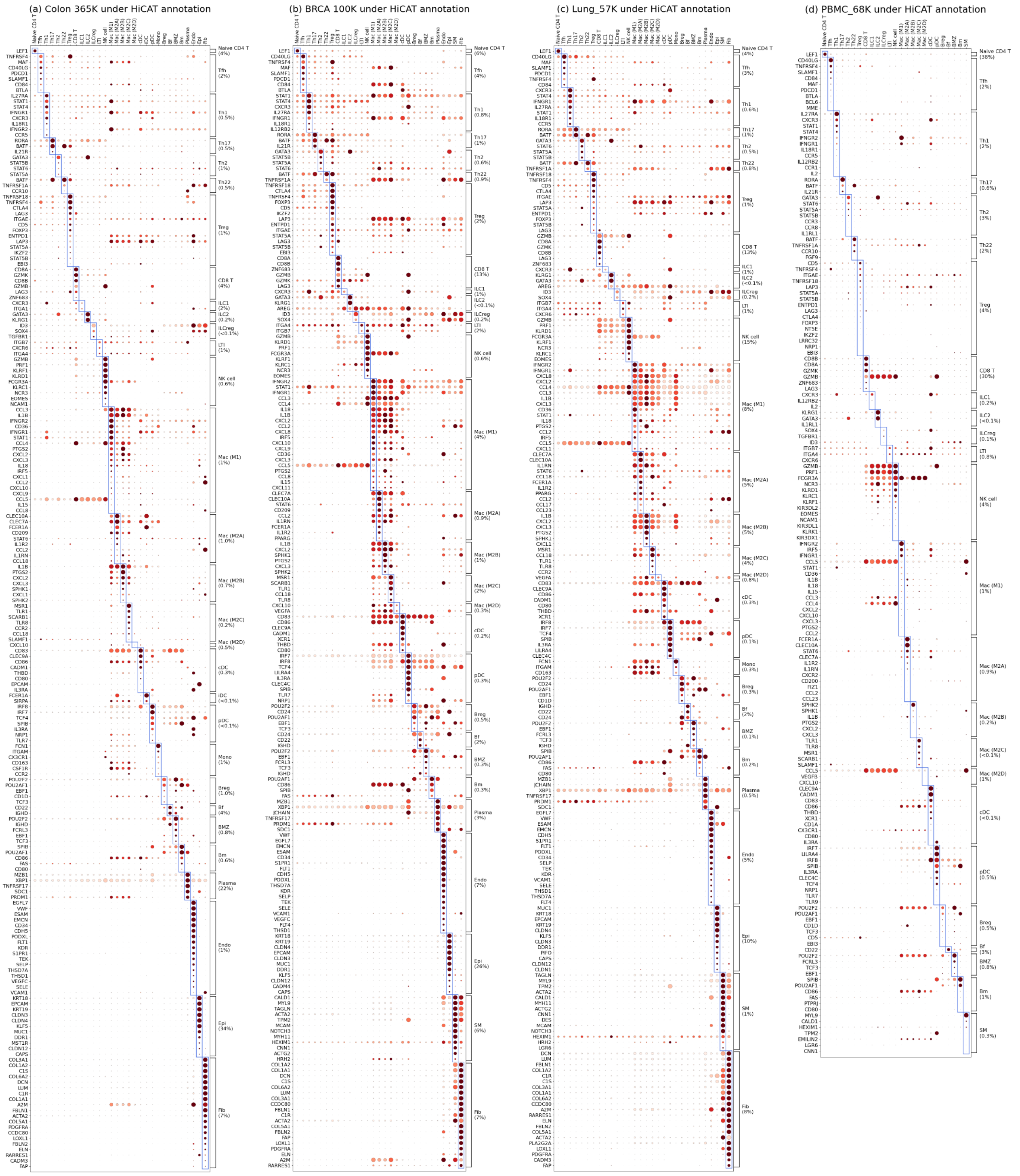


**Supplemental Figure 5**. Dot plots showing marker expression profile under HiCAT annotation in (a) Colon 365K, (b) BRCA 100K, (c) Lung 57K, and (d) PBMC 8K and. The dot size is the fraction of cells that expresses the marker gene in the identified subsets, regardless of their expression level. The color corresponds to the average expression level of the marker scaled between 0 and 1, where 0 and 1 represent respectively the minimum and the maximum expression level of that marker among all subsets. The fraction was clipped at 0.6. For better visualization, we grouped the marker genes for each subset and ordered according to the fraction of marker expressing cells in the designated subset. We removed some common marker that frequently occurs (in 3 or more subsets), e.g., *CD3* and *CD4* in T cell subset, *CD14* in myeloid cell subsets, *PTPRC*, *IL7R* and *KLRB1* in ILC subsets, *IGHM*, *CD19*, *CD27*, *MS4A1* in B cell subsets. We showed up to 180 markers with highest percentage of marker expressing cells.

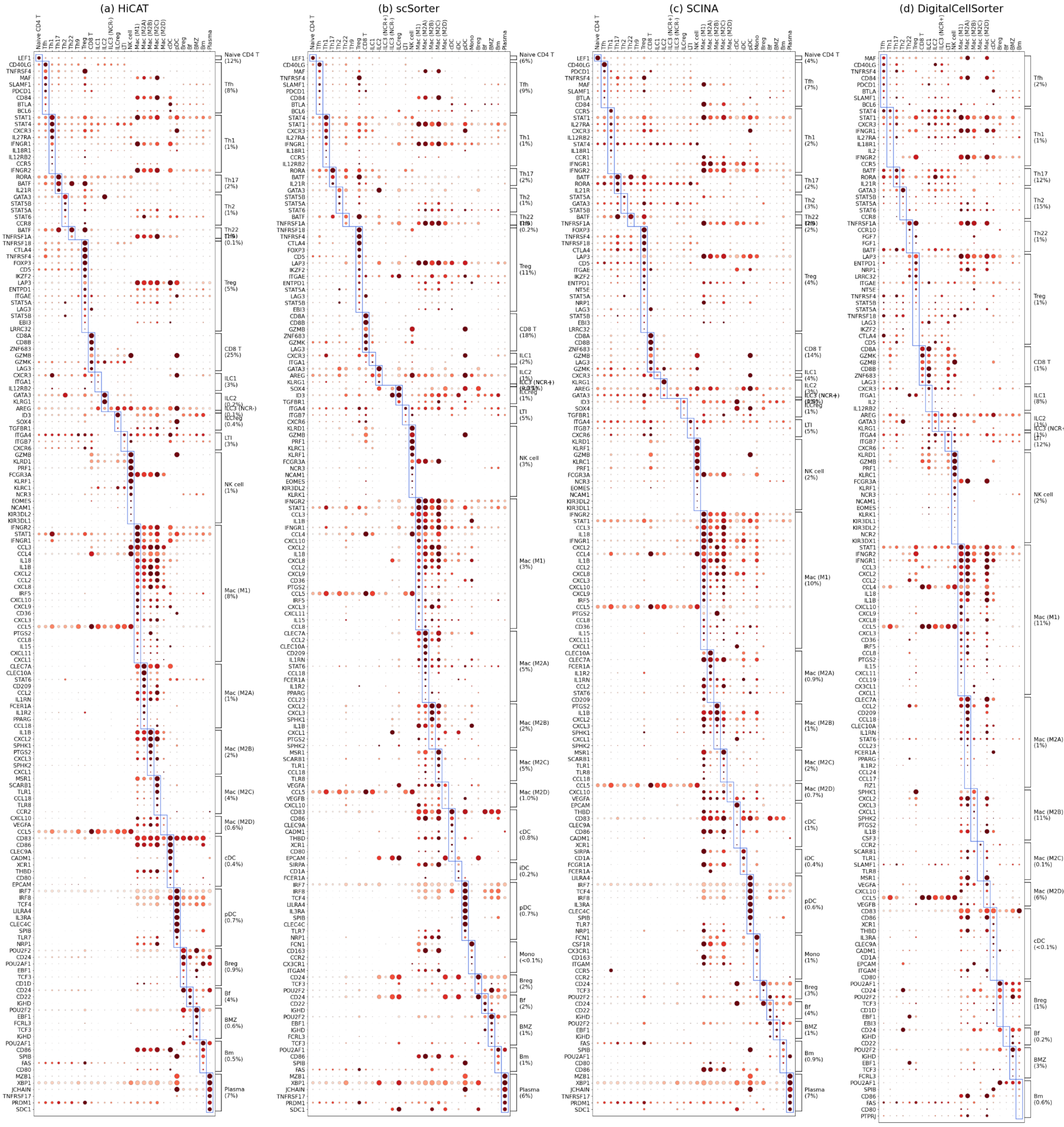


**Supplemental Figure 6**. Dot plots showing marker expression of immune cell subsets in BRCA 100K identified by different tools. The dot size is the fraction of cells in an identified subset that expresses the marker, regardless of their expression level. The color corresponds to the expression level scaled between 0 and 1, where 0 and 1 represent respectively the minimum and the maximum expression level of that marker among all subsets. The fraction was clipped at 0.6, i.e., fractions greater than 0.6 were set to 0.6. For better visualization, we grouped the marker genes for each subset and ordered according to the fraction of marker expressing cells in the target subsets. Each column is a subset identified by each method. We removed some common marker that frequently occurs (in 3 or more subsets), e.g., *CD3* and *CD4* in T cell subset, *CD14* in myeloid cell subsets, *PTPRC*, *IL7R* and *KLRB1* in ILC subsets, *IGHM*, *CD19*, *CD27*, *MS4A1* in B cell subsets. We showed up to 180 markers with highest percentage of marker expressing cells.
